# Different Effects of Three Types of Water on Developmental Behaviors, Lipid Metabolism and Antioxidant Capacity of Juvenile Zebrafish

**DOI:** 10.1101/373381

**Authors:** Cheng Chang, Jiajin Zhu

## Abstract

Adolescence is an important period when people need adequate nutrition for healthy growth and development, whether from food or water. Tap water, bottled water (bottled purified water, and bottled natural water) are now the three most popular drinking waters consumed by adolescents in China. However, the constituents of them differ, which may cause different long-term health effects. In order to determine which type of water is the most beneficial regarding developmental behaviors, lipid metabolism and antioxidant capacity of juvenile zebrafish, 21 dpf (days post-fertilization) zebrafish were given these three waters separately with the same feed until 90 dpf. Results showed that zebrafish in purified water had the lowest survival rate, body weight and body length, while zebrafish in natural water and tap water had similar developmental behaviors; the highest HDL-C level and the lowest TG level were found in natural water, tap water second; and zebrafish in natural water showed the best antioxidant capacity. Thus, the best outcomes were found in natural water, which had suitable pH and proper amount of minerals, tap water also showed good performance, while purified water seemed not suitable for juvenile zebrafish.

## Introduction

Adolescence is an important period for growth and development, which determines the physique, intelligence and spirit state of human body (Das et al., 2017). Whether adolescents can grow up healthily is not only influenced by genetic factors, but also closely related to proper nutrition, particularly various macro and trace mineral elements. Due to unreasonable dietary structure of the Chinese residents, the deficiency of calcium, zinc, selenium and magnesium is serious (Fan et al., 2012). The survey shows that the dietary calcium intake of Chinese residents is generally on the low side. Among them, young people aged 11 to 13 reaches the lowest proportion of DRIs (Chinese Dietary Guide). Similarly, the survey data from the United States also show that the ratio of dietary calcium intake to AI (Adequate Intakes) is the lowest among children and adolescents aged 8 to 19. At the same time, due to increased dependence on over-refined staple foods such as cereals and carbohydrates, dietary magnesium intake is no longer able to meet recommended intakes.

The human body consumes mineral elements mainly through eating food and drinking water. Compared with food, mineral elements in water are mostly ionized and therefore more easily absorbed. Sabatier et al. suggested that the bioavailability of magnesium and calcium in water is higher than that in food (60% vs 40%) (Sabatier, Arnaud, Kastenmayer, Rytz, & Barclay, 2002). Due to the particularity of adolescents, they actually drink boiled water and bottled water daily. As raw material for boiled water, tap water supplies 10% of the average individual’s zinc intake. However, tap water is susceptible to biological and chemical pollution (Cidu, Frau, & Tore, 2011), during the process of tap water disinfection, all kinds of disinfectants may produce disinfection by-products (DBPs) in water, especially trichloromethane (Richardson (2003);Richardson, Plewa, Wagner, Schoeny, & DeMarini, 2007). In addition, copper, a heavy metal element, can be leached from pipes and faucets into the tap water stream (Cartier, Nour, Richer, Deshommes, & Prevost, 2012). Therefore, unpleasant odors from disinfectants, DBPs, and heavy metal exposure can drive people to choose bottled drinking water increasingly. The most popular bottled drinking water in the market is bottled natural water and bottled purified water. Due to differences in water sources and filtration methods, bottled purified water contains almost no mineral elements, while bottled natural water contains a certain amount of mineral elements. Therefore, we can see that different bottled water contains different levels of mineral elements, but they are all lower than that in tap water.

The three popular drinking water have different contents of components and elements, which may lead to different effects on body health. Studies have found that different types of drinking water have an impact on skeletal development, lipid metabolism and antioxidant capacity. Long-term consumption of purified water may have negative effects on lipoprotein metabolism (Zhao, Shu, & Gao, 2004). Ling et al. found that zinc addition in water increased lipolysis and decreased lipogenesis of goby Synechogobius hasta (Ling, Luo, Chen, Zhang, & Liu, 2018). Furthermore, the mortality of coronary heart disease or other cardiovascular disease in soft water area is higher than that in hard water area (Nerbrand, Agreus, Lenner, Nyberg, & Svardsudd, 2003). Studies have shown that long-term drinking soft water can lead to insufficient magnesium intake and lipid peroxidation. All these studies remind us to pay attention to the choice of drinking water in our daily life, especially the adolescents who are in the stage of growth and development. However, few studies have compared the effects of long-term drinking of different drinking water on adolescents. What’s more, it is well-recognized that water which contain mineral elements are more beneficial for human being, but in this experiment, we concentrated on studying whether mineral elements in water still show positive effects when body can get adequate mineral elements from food.

Zebrafish (*Danio Rerio*), as a new type of vertebrate whose gene is up to 87% homologous to humans (Tsang, 2010), has many advantages that other traditional model animals do not have, such as small size, short life cycle and easy feeding. In addition, zebrafish is an in vitro fertilized and in vitro developed animal whose physiological process is highly similar to that of human beings (Gerlach, Morales, & Wingert, 2015), making it popular among scientific researchers. The water environment has great influence on the growth, development, physiology and pathology of zebrafish (Oberemm, 2000), therefore, the International Standards Organization (ISO) recommended zebrafish as experimental species for testing river water toxicity, and formulated a corresponding standard (ISO07346). China has also introduced the national standard GB/T 13267-1991 (Water Quality - Determination of the acute toxicity substances of a freshwater fish) with reference to international standards. The use of zebrafish for water quality safety has been recognized and widely used in China and abroad. In this study, we compared the effects of three popular drinking water on development, lipid metabolism and antioxidant capacity of juvenile zebrafish.

## Materials and methods

### Animals

All animal procedures were performed in full accordance to the requirement by “Regulation for the Use of Experimental Animals in Zhejiang Province”. This work is approved by the Animal Ethics Committee in the School of Medicine, Zhejiang University (Ethics Code Permit No. ZJU2015-8-26-004Y, issued by the Animal Ethics Committee in the School of Medicine).

Zebrafish (Danio rerio) were raised and maintained in the standard Zebrafish Unit (produced by Aisheng Zebrafish Facility Manufacturer Company, Beijing, China) at Zhejiang University under a constant 14 h on/10 h off light cycle at 28°C. Wild type AB strain zebrafish embryos were obtained from the Key Laboratory for Molecular Animal Nutrition, Ministry of Education, College of Animal Sciences, Zhejiang University (Hangzhou, China) and were used in this study. The embryos were raised in standard E3 solution without methylene blue, at 28°C and on a 14/10 light/dark cycle^[24]^. At 5 dpf (days post-fertilization), the larvae fish were moved to fish water and fed twice a day until 21 dpf. At the end of the experiment, the zebrafish were euthanised using MS222 (Sigma-Aldrich).

### Water and diet

The tap water used in the experiment was municipal water of Hangzhou, after 24 hours of aeration; The bottled natural water and bottled purified water were purchased from the supermarket; The fish water was made up of purified water with sea salt, with the electrical conductivity adjusted to 290 ~ 320 us/ cm.

The zebrafish feed was Zeigler larval (crude protein > 50.0%, crude fat > 12.0%, crude fibre<2.5%).

### Instruments and Equipment

Atomic Absorption Spectroscopy (AAS), AA240, Varian; Inductively coupled plasma emission spectrometer (ICPAES), 730-ES, Varian; pH meter, METTLER TOLEDO; Digital blower dryer, GZX-9030MBE, BOXUN; Intelligent biochemical incubator, SPX; Double beam ultraviolet visible spectrophotometer, TU-1900, PERSEE.

### Kits

All the kits were bought from Nanjing Jiancheng Bioengineering Institute. Total protein quantitative assay kit, A045-2; Triglyceride assay kit, A110-2; Total cholesterol assay kit, A111-2; High-density lipoprotein cholesterol assay kit, A112-2; Low-density lipoprotein cholesterol assay kit, A113-2; Malondialdehyde (MDA) assay kit, A003-1; Total Superoxide Dismutase (T-SOD) assay kit, A001-1-1.

### Determination of quality and components of water

The pH, total dissolved solids (TDS), and potassium, calcium, sodium, magnesium, iron, zinc, copper levels were measured according to GB5749-2006. Among them, the pH value was determined using the glass electrode method; TDS was determined by the weighing method, with the drying temperature 105°C ±3°C; the potassium level was determined by flame atomic absorption spectrometry; the calcium, sodium, magnesium, iron, zinc, and copper levels were determined by ICPAES.

### Intervention procedure

21 dpf juvenile zebrafish were randomly divided into 4 groups: TWG (fish water group) (control group), PWG (purified water group), NWG (natural water group) and TWG (tap water group), 50 tails in each group, as shown in Figure 1. Each group was raised in the corresponding water, and 1/3 of the fresh water was changed daily to ensure the oxygen content in the water. All the juvenile zebrafish were fed with the same feed during the experiments to ensure that the nutrient contents of the feed had no effect on the experimental results. The zebrafish freely ate and the amount of feed was guaranteed to be eaten within 5 min each time. The zebrafish were feed 2 times a day early and late.

**Figure 1.**
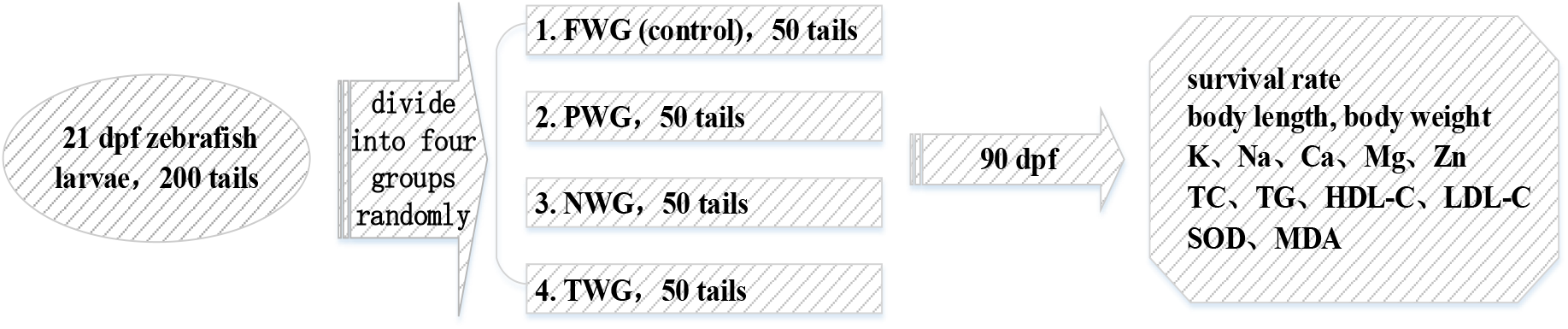
Schematic illustration of the study design

### Survival rate

During the experiments, the number of zebrafish deaths in every group was recorded daily and the survival rate was plotted.

### Body Weight and Body length

24 h before the end of the experiment, all zebrafish were stopped feeding. On experimental day, the zebrafish were euthanised using MS222 (Sigma-Aldrich) and the residual water on the surface was absorbed by filter paper thoroughly, then the body weight and body length of the whole fish were measured.

### K, Na, Ca, Mg, Zn, Fe, Cu levels

After the zebrafish were euthanised, 9 volumes of normal saline were added to the zebrafish in a ratio of weight (g): volume (mL)=1:9, and the fish was cut with a surgical cutter and homogenized with an ultrasonic crusher in an ice water bath. Until homogenization of the tissue mixture, mixing, centrifugation at 4°C, 5000×g for 30 min, and taking the supernatant to obtain a 10% zebrafish homogenate. After diluted 10 times, K level of the whole fish was determined by flame atomic absorption spectrometry. Na, Ca, Mg, Zn, Fe, Cu levels were determined by inductively coupled plasma atomic emission spectrometry.

### Lipid Metabolism

The contents of protein, TG, TC, HDL-C and LDL-C were determined according to the corresponding kit instructions.

### Antioxidant Capacity

SOD activity and MDA level were measured according to the corresponding kit instructions.

### Statistical analysis

All statistical analyses were performed using SPSS 16.0 Statistical Software. Data were analyzed using one-way ANOVA followed by a post hoc LSD test. The Pearson bivariate correlation analysis test was performed to analyze the relationship between the levels of minerals and lipid metabolism or anti-oxidation parameters. All the data were presented as mean ± SEM. A p value smaller than 0.05 was considered to be statistically significant.

## Results

### Quality and constituents of the water

Table 1 shows that compared with tap water, the two bottled water have lower pH and contained relatively lower levels of total dissolved solids (TDS), indicating that the levels of inorganic components were lower in the bottled waters. Macro elements such as potassium, sodium, calcium, and magnesium were higher in the tap water than in the two bottled waters, and among the two types of bottled water, these macro elements were higher in natural water than in purified water. Furthermore, the calcium and magnesium ratio was 13:1 in purified water but 3:1 in fish water, 7:1 in natural water and 8:1 in tap water. Zinc, another important component, was highest in tap water as expected. Besides, iron and copper are trace in all four types of water, far below the limits specified in GB 5749-2006 ( iron<0.3 and copper<1.0).

**Table 1.** The water quality indexes and components of the four waters.

|  | **Fish Water** | **Purified Water** | **Natural Water** | **Tap Water** | **Unit** |
| --- | --- | --- | --- | --- | --- |
| **pH** | 7.14 | 6.78 | 7.24 | 7.60 |  |
| **TDS** | 204 | 5 | 50 | 165 | mg/L |
| **K** | 0.6747 | 0.1503 | 1.0742 | 2.9738 | mg/L |
| **Na** | 85.1273 | 0.3022 | 3.8981 | 16.5123 | mg/L |
| **Ca** | 2.3974 | 0.1192 | 12.8940 | 27.4650 | mg/L |
| **Mg** | 0.6988 | 0.0093 | 1.8659 | 3.4035 | mg/L |
| **Zn** | 0.0555 | 0.0341 | 0.0344 | 0.2755 | mg/L |
| **Fe** | 0.0209 | 0.0208 | 0.0209 | 0.0052 | mg/L |
| **Cu** | 0.0015 | 0.0017 | 0.0023 | 0.0003 | mg/L |

### Developmental parameters of juvenile zebrafish

21 dpf, which was the first day of the experiment, there have been several deaths of juvenile zebrafish in PWG and NWG, leading to lower survival rate than that in FWG and TWG(Fig. 2). 25 dpf, juvenile zebrafish in PWG continued to die in large amount, and juvenile zebrafish in TWG began to die, while mortality in FWG and NWG flattened out. 60 dpf, juvenile zebrafish in NWG, TWG and FWG all stopped the death, while there were still a few deaths in PWG. 90 dpf, the survival rate in PWG was the lowest among the four groups, survival rates in the other three groups were similar.

**Figure 2.**
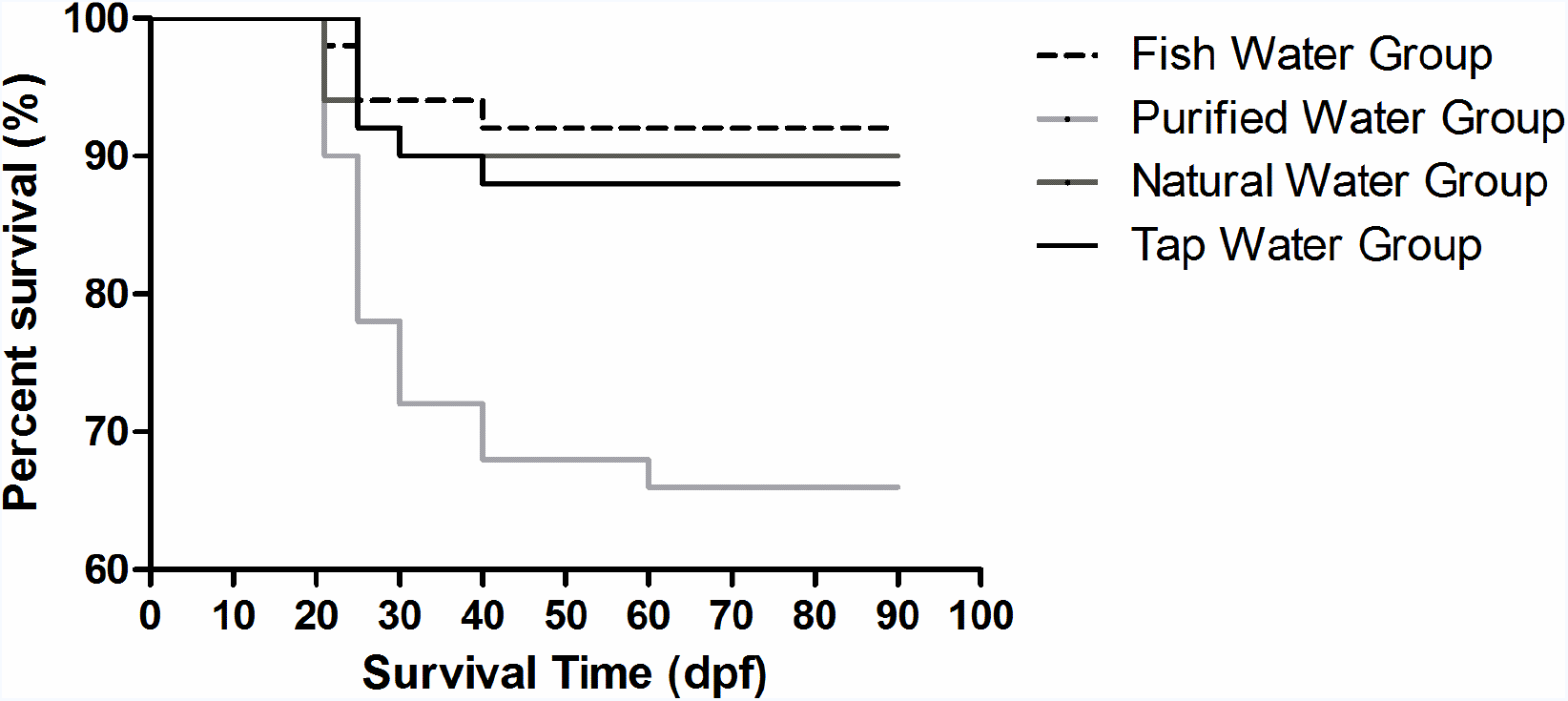
The survival curve of zebrafish juvenile in four groups.

Table 2 showed that the Body Weight and Body Length of 90 dpf zebrafish in PWG was significantly lower than that in NWG (p=0.023, 0.002 separately), but there was no difference of that in 21 dpf zebrafish. No significant differences of Body Weight and Body Length were observed in other groups. What’s more, no significant difference was observed in BMI, but there is a decreasing tendency in NWG.

**Table 2.**
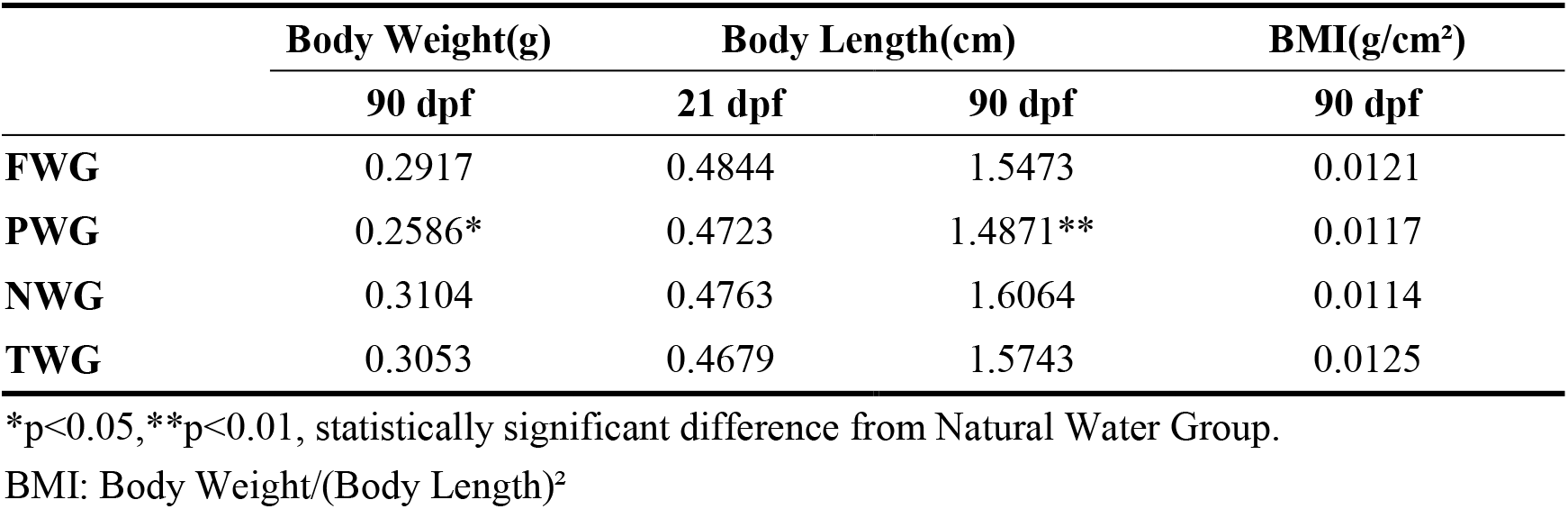
The body weight and body length of 90 dpf zebrafish.

### Potassium, sodium, calcium, magnesium, iron, copper and zinc levels in 90 dpf zebrafish after experiment

There were no significant differences in mineral levels of potassium, sodium, magnesium and copper among all four groups (Fig. 3A,B,D,G). Calcium level in NWG was significantly higher than that in PWG and TWG (p=0.013, 0.022 separately) (Fig. 3C). Iron level in FWG was significantly higher than that in all other three groups, and the level in PWG was significantly higher than that in TWG (Fig. 3E). Furthermore, zinc level in FWG was significantly lower than NWG and TWG, and zinc level in PWG was the lowest among all the four groups (Fig. 3F).

**Figure 3.**
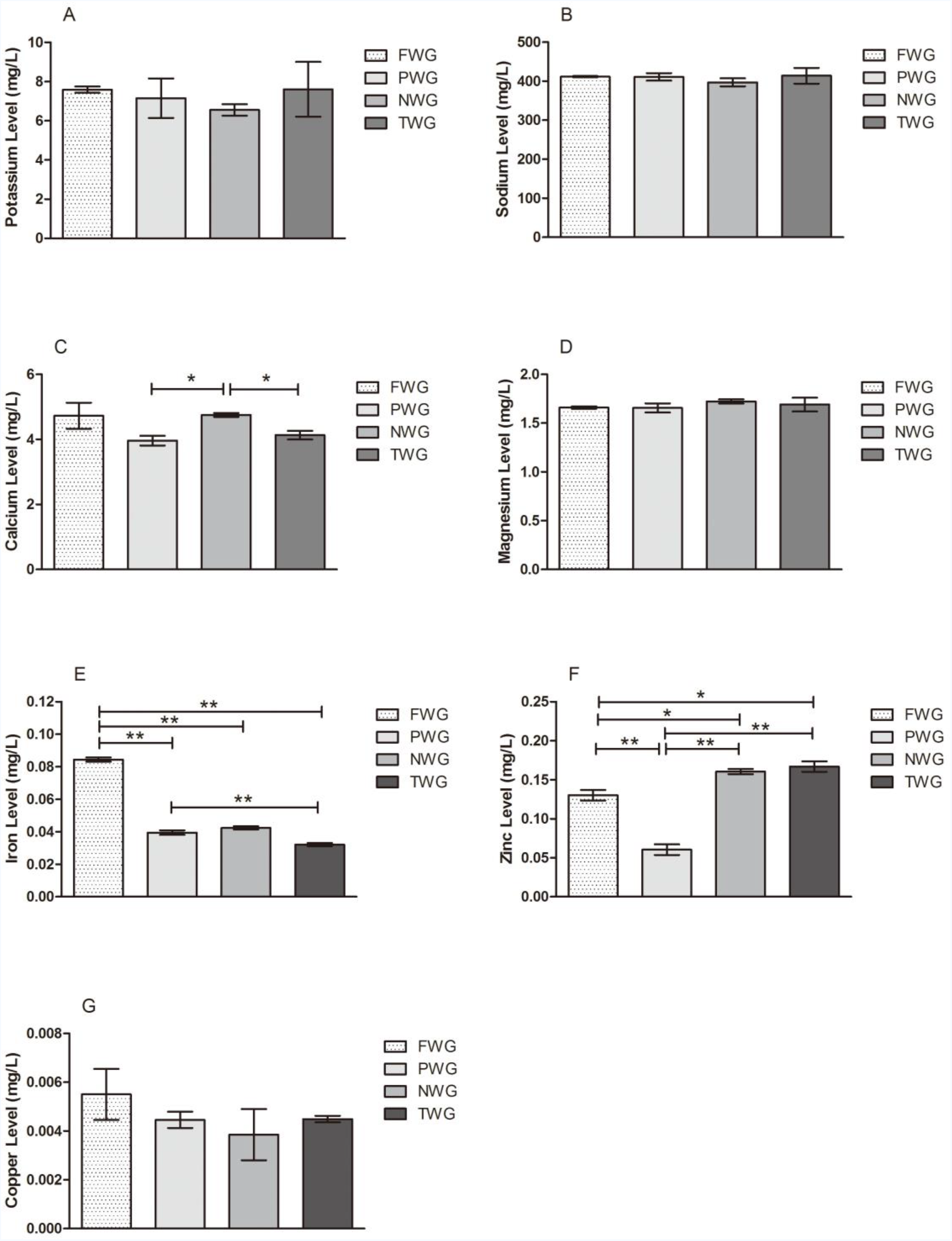
The mineral levels of 90 dpf zebrafish in four different waters ( ̄±s). Statistically significant differences are marked with asterisks: *p<0.05 and **p<0.01.

### 90 dpf zebrafish lipid metabolism

After the experiment, TG level of zebrafish in TWG dropped compared with FWG, and TG level in PWG was significant higher than NWG (p=0.016) and TWG (p=0.001) (Fig. 4A). What’s more, zebrafish in NWG exhibited statistically significantly higher HDL-C level than FWG and PWG (p=0.002, 0.041 separately) (C). There were no significant differences in TC level and LDL-C level between the four groups.

**Figure 4.**
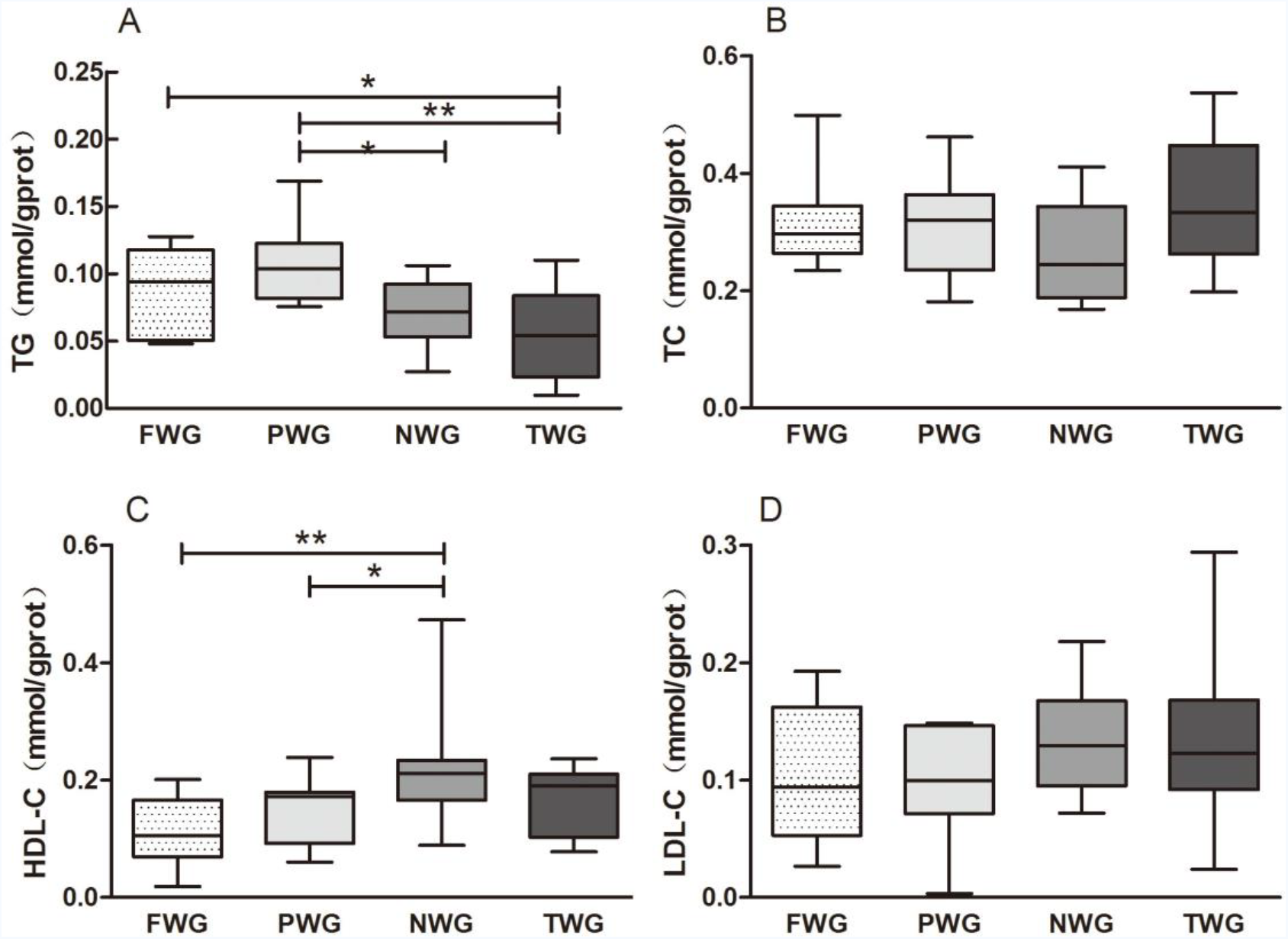
90 dpf zebrafish TG level(A), TC level(B), HDL-C level(C) and LDL-C level(D). Statistically significant differences are marked with asterisks: *p<0.05 and **p<0.01.

### 90 dpf zebrafish antioxidant capacity

There were no significant differences in SOD activity of 90 dpf zebrafish after the experiment between all four groups (Fig. 5A). However, MDA content in PWG was significantly higher than that in NWG (Fig. 5b).

**Figure 5.**
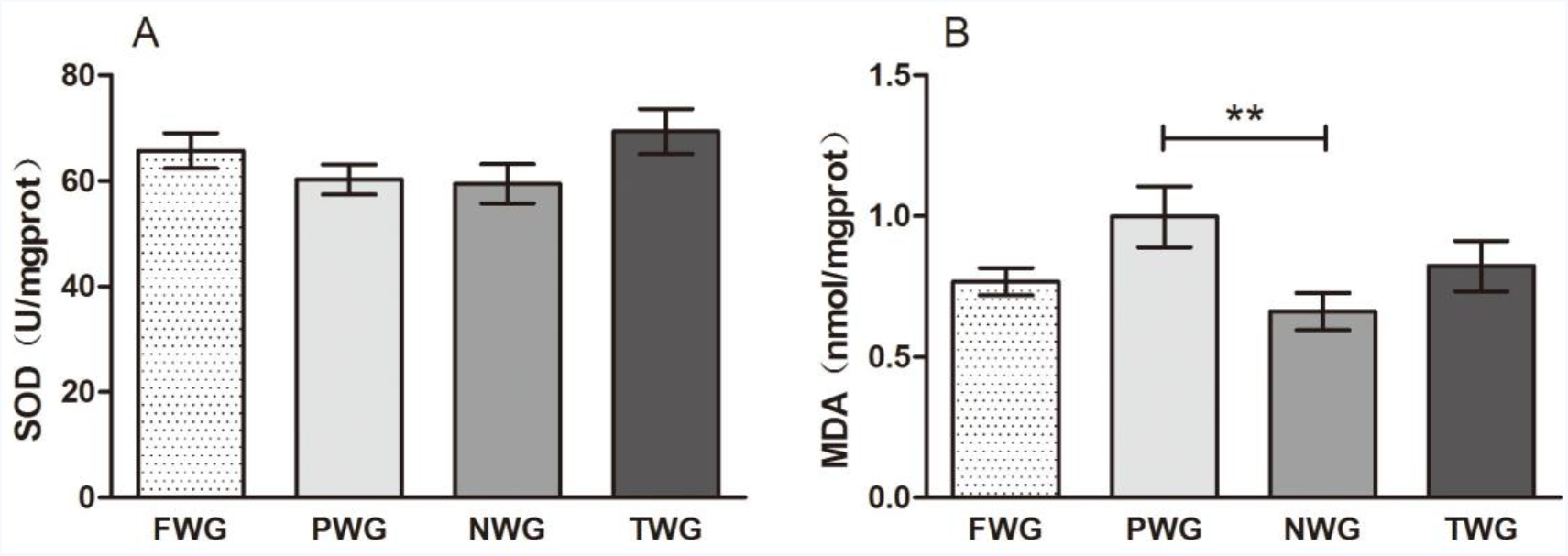
90 dpf zebrafish SOD(A) and MDA (B). Statistically significant differences are marked with asterisks: **p<0.01.

### The relationship of water constituents, zebrafish developmental parameters, mineral levels, lipid metabolism and antioxidant parameters

Table 3 showed that survival rate of zebrafish was associated with all the water constituents, zebrafish body length was positively associated with water calcium (p=0.04) and magnesium levels (p=0.02). We can also notice that TG level of zebrafish was positively associated with water potassium level (p=0.001), but was negatively associated with water calcium (p=0.001), magnesium (p=0.000), iron (p=0.01) and zinc levels (p=0.01). Furthermore, HDL-C level of zebrafish was negatively associated with water sodium level (p=0.022) and negatively associated with water calcium and magnesium levels.

**Table 3.** Correlation analysis of the relationship among water constituents, zebrafish developmental parameters, mineral levels, lipid metabolism and antioxidant parameters.

|  | **Survival**<br>**Rate** | **Body**<br>**Weight** | **Body**<br>**Length** | **Zebrafish**<br>**TG** | **Zebrafish**<br>**TC** | **Zebrafish**<br>**HDL-C** | **Zebrafish**<br>**LDL-C** |
| --- | --- | --- | --- | --- | --- | --- | --- |
| **Water pH** | 0.733** | 0.286* | 0.321* | 0.539** | 0.134 | 0.136 | 0.228 |
| **Water TDS** | 0.860** | 0.167 | 0.127 | -0.221 | 0.192 | -0.279 | 0.053 |
| **Water Potassium** | 0.5** | 0.224 | 0.235 | 0.505** | 0.203 | 0.118 | 0.203 |
| **Water Sodium** | 0.64** | 0.039 | -0.004 | 0.085 | 0.075 | -0.374* | -0.076 |
| **Water Calcium** | 0.434** | 0.239 | 0.273* | -0.522** | 0.141 | 0.229 | 0.231 |
| **Water Magnesium** | 0.538** | 0.265 | 0.307* | -0.54** | 0.122 | 0.227 | 0.241 |
| **Water Iron** | -0.293* | -0.142 | -0.113 | -0.414** | -0.277 | -0.021 | -0.144 |
| **Water Zinc** | 0.356** | 0.147 | 0.115 | -0.413* | 0.286 | -0.014 | 0.139 |
| **Water Copper** | -0.275* | -0.064 | 0.014 | 0.274 | -0.349* | 0.198 | -0.051 |
| **Zebrafish Sodium** | -0.032 | -0.793** | -0.605* | -0.6 | -0.115 | -0.011* | -0.163 |
| **Zebrafish Calcium** | 0.452* | -0.068 | 0.109 | -0.37 | -0.554** | 0.185 | -0.286 |
| **Zebrafish Magnesium** | 0.168 | -0.59 | -0.255 | -0.677* | -0.33 | 0.483 | 0.268 |
| **Zebrafish Zinc** | 0.839** | 0.768* | 0.761* | -0.218 | -0.111 | 0.794* | 0.702 |

|  | **Zebrafish**<br>**SOD** | **Zebrafish**<br>**MDA** | **Zebrafish**<br>**Calcium** | **Zebrafish**<br>**Magnesium** | **Zebrafish**<br>**Iron** | **Zebrafish**<br>**Zinc** |
| --- | --- | --- | --- | --- | --- | --- |
| **Water pH** | 0.255 | -0.233 | 0.132 | 0.211 | -0.214 | 0.907** |
| **Water TDS** | 0.3 | -0.147 | 0.191 | -0.027 | 0.57 | 0.542 |
| **Water Potassium** | 0.269 | -0.103 | -0.08 | 0.154 | -0.433 | 0.741* |
| **Water Sodium** | 0.157 | -0.104 | 0.312 | -0.143 | 0.936** | 0.132 |
| **Water Calcium** | 0.212 | -0.139 | -0.039 | 0.216 | -0.554 | 0.771* |
| **Water Magnesium** | 0.216 | -0.188 | 0.041 | 0.234 | -0.462 | 0.843** |
| **Water Iron** | -0.285 | -0.036 | 0.277 | -0.051 | 0.495 | -0.502 |
| **Water Zinc** | 0.302 | 0.026 | -0.252 | 0.041 | -0.417 | 0.522 |
| **Water Copper** | -0.327* | -0.133 | 0.353 | 0.09 | 0.222 | -0.275 |
| **Zebrafish Sodium** | -0.351 | 0.082 | 0.187 | 0.653* | 0.005 | -0.799* |
| **Zebrafish Calcium** | -0.295 | -0.249 | 1 | 0.583 | 0.19 | 0.145 |
| **Zebrafish Magnesium** | -0.111 | 0.098 | 0.583 | 1 | -0.132 | -0.206 |
| **Zebrafish Zinc** | 0.344 | 0.161 | 0.145 | -0.206 | -0.055 | 1 |

\*, p<0.05; \*\*, p<0.01.

## Discussion

### Water components

As is known, different water source and treatment procedures may result in different water constituents. From our study, the data showed different pH and TDS levels in the three popular types of water, furthermore, the TDS levels were lower in two bottled water than tap water, which indicated that after filtration treatment, the majority of the minerals were removed. According to R.J. Erickson unpublished data, the Ca:Mg ratio in natural waters ranges from 1.6:1 to 8:1 (Naddy, Stubblefield, May, Tucker, & Hockett, 2002). Our data showed that apart from bottled purified water, Ca:Mg ratio in the other two types of water were within this range, which means that more magnesium than calcium were removed after bottled purified water being filtered. K, Na, Ca, Mg, Zn levels in tap water were much higher than those in two types of bottled water, and more in bottled natural water than in bottled purified water, which is consistent with the theoretical value and several studies (Carstea, Levei, Hoaghia, & Savastru (2016);Zeng et al., 2014). What’s more, our study showed that copper and iron levels of tap water were in accordance with GB5749-2006 in China. It is worth noting that TDS in fish water was highest among the four waters, this may be due to the fact that the sodium content in fish water is much higher than the other three types of water.

### Effects of water constituents on juvenile zebrafish development

A previous study have suggested that fish are most suitable developed in neutral or alkalescent waters, living in acidic environment will lead to inactivity, low metabolic rate, and reduced food intake. Lawrence et al. approved that while most freshwater fish can tolerate a wider pH range of 6.0~9.5, it is generally practical to maintain them at a pH in the 7.0~8.0 range in order to promote good health of biofilters and stable water quality (Lawrence, 2007). In general, zebrafish larvae have stricter requirements than adults and should be raised in egg water with pH adjusted to approximately 7.2. That is to say, apart from bottled purified water, all the water in our experiment were suitable for zebrafish juveniles to develop, which is consistent with the survival rate.

Zhang et al. reported that drinking purified water can significantly reduce body weight of rat when taking magnesium deficient diet (Zhang, Fan, & Gu, 2007). Several evidence suggested that calcium intake before puberty influences adolescent height growth (Bhargava, 2016); Prentice, Dibba, Sawo, & Cole, 2012), calcium intake below 300 mg/d may result in shorter adult stature (Fang et al., 2017). Our study showed that body weight and body length of juvenile zebrafish decreased significantly in purified water group, compared with the other three groups. The correlation analysis also showed that the juveniles’ body length was positively correlated to calcium and magnesium levels in water. The juveniles’ body weight was positively correlated to its zinc levels, and zinc levels in juvenile zebrafish were positively correlated to calcium and magnesium levels in water. Thus, the significantly decreased weight and body length in purified water group might due to the deficiency of calcium and magnesium in purified water.

### Effects of water constituents on juvenile zebrafish lipid metabolism

Coronary artery disease is a major cause of morbidity and has various risk factors. Lipid profile i.e. low HDL-cholesterol, high LDL-cholesterol, high total cholesterol, high triglycerides playing important role in its causation (Ghosh, Mishra, Rao, & Aggarwal, 2006). The role of triglycerides in atherosclerosis had been neglected for many years as it was a widely held view that elevated triglycerides do not represent a risk factor for this disease. However, It is known that increased hepatic synthesis and production of TG are frequently associated with insulin resistance, and these metabolic disturbances often lead to TG accumulation in the liver, contributing to hypertriglyceridemia (Bacon, Farahvash, Janney, & Neuschwandertetri, 1994). The correlation analysis showed that the juveniles’ TG level was negatively correlated to calcium, magnesium, iron, zinc levels and positively correlated to potassium level in water. Papakonstantinou et al. found that rats fed high-calcium diet had 29% less carcass fat than controls (Papakonstantinou, Flatt, Huth, & Harris, 2003). It is well established that high concentrations of calcium in the diet can inhibit absorption of dietary fatty acids, suppressing circulating concentrations of hormones such as 25-hydroxyvitamin D, 1,25-dihydroxyvitamin D, and parathyroid hormone, which normally promote calcium uptake and influx into cells, and thus, reverse the adipogenic action of Agouti protein by reducing intracellular calcium concentrations (Zemel, Shi, Greer, Dirienzo, &Zemel, 2000). Furthermore, Hanne et al. found that magnesium significantly reduced TG levels in apoE^-/-^ mice when the animals received standard mouse chow with a sufficient magnesium content (Ravn, Korsholm, & Falk, 2001). However, Orimo and Ouchi found unaltered lipid levels in cholesterol-fed rabbits (Orimo & Ouchi, 1990). The reason for the discrepancy between the two studies might be related to the fact that magnesium was administered in the drinking water in the first study but as part of the diet in the latter study. It has been argued that the bioavailability of magnesium is higher from water than from food, which in theory would also contribute to a reduced effect. What’s more, since the iron content in the three waters is similar and trace, we can infer that the iron level range in our experiment didn’t result in changes in zebrafish TG levels, so do zinc. Therefore, we can say that it is the difference of calcium and magnesium levels in water that causes the change of TG level.

The cardioprotective effect of HDL has been largely attributed to its role in reverse cholesterol transport (RCT), wherein excessive cholesterol is conveyed from peripheral tissues to the liver and steroidogenic organs (Tian et al., 2010). The correlation analysis showed that the juveniles’ HDL-C level was positively correlated to calcium, magnesium and negatively correlated to sodium level in water. The results were in consistent with other studies: Delisle et al. found that intakes of calcium were significantly better in adults with normal HDL-C level than those with low HDL-C level (Delisle, Ntandou, Sodjinou, Couillard, & Despres, 2013), and HDL-C was significantly higher with increasing quartile of magnesium intake (Jui-Hua, Yi-Fa, Fu-Chou, Ning-Yuean Lee, & Leih-Ching, 2012). Choi et al. presented that dietary sodium intake and serum HDL-C had a negative correlation (r=-0.11, p<0.05) (Choi, Lee, & Park, 2005). However, there are some studies that show inconsistent results: Aslanabadi et al. found that one-month intake of mineral water rich in calcium, magnesium bicarbonate, and sulfate decreased cholesterol and LDL levels but not TG or HDL levels in dyslipidemic adults (Aslanabadi, Habibi Asl, Bakhshalizadeh, Ghaderi, & Nemati, 2014). We hypothesized that it might be due to the water used in the control group still contained higher concentrations of calcium (7 ppm) and magnesium (1.7 ppm) than our experiment.

Mineral-rich water could provide an important supplementary contribution to total calcium and magnesium intake according to the results of a French study (Mooradian, 2009). The suggested mechanisms are related to the moderately alkaline nature of the study mineral water and an osmotic effect that may affect fat and cholesterol absorption and/or increase bile acid excretion. It is known that the rate of fatty acid and cholesterol absorption from the micellar solution formed in the small intestine desires a lower pH and that the action of pancreatic enzymes and bile salts is increased by the addition of pH. Therefore, an increase in luminal pH induced by the mineral water may decrease the uptake of both cholesterol and fat. Various mineral waters are able to increase the excretion of bile consumed with or without a meal (Albertini M C, 2007). So we can concluded that the significant changes of TG and HDL-C levels in the purified water group were caused by the lack of calcium and magnesium and lower pH level in bottled purified water.

### Effects of water constituents on juvenile zebrafish antioxidant capacity

To defend themselves against oxidative damage from free radicals, organisms start an emergency response mechanism, such as increased activity of antioxidant enzymes SOD, CAT and GPx, and reduced endogenous antioxidant (GSH) (Xing et al., 2012). Superoxide dismutase (SOD; an enzyme that catalyzes the dismutation of the superoxide radical into oxygen or hydrogen peroxide) represent the first line of cellular antioxidant protection against oxidative damage caused by ROS and lipid peroxidation (Wang, Zhong, Shi, & Guo, 2011), and fish are inferred to degrade superoxide into oxygen using SOD (Khan R A, 2012). Interestingly, SOD activity of juvenile zebrafish wasn’t significantly affected by different waters, indicating that all the zebrafish exhibited similar antioxidant capacity. Our results were in contrast with what was found for other seawater animals (Pimentel et al., 2015);Rosa et al., 2014), but were consistent with subfreshwater fishes. The reasons could be that freshwater fish are usually exposed to higher pH variations (daily and throughout the year) than seawater animals and thus the change of 0.8 units on pH was not enough to induce an oxidative stress response. Moreover, it has been reported that magnesium has an anti-lipid peroxidation effect, and can maintain the activity of SOD. Han et al. found that when supplemented with 1 mg/kg magnesium, SOD activities markedly increased in high fluoride cattle (Han et al., 2004). This is inconsistent with our results, probably because there was only small difference in magnesium levels in our experiment, and our zebrafish were not in oxidative stress.

Once free radicals breaks through the organisms’ defenses against oxidation, ROS may attack cellular molecules that induce oxidative damage, thus we assessed changes in lipid peroxidation in juvenile zebrafish by measuring MDA content. MDA content is considered to be one of the most important manifestations of lipid peroxidation (Hopps, Noto, Caimi, & Averna, 2010). Our result was consistent with Wei-qun Shu et al, the increased MDA levels in purified water group in our study may be attributed to the deficiency of magnesium. It is reported that magnesium has the effect of anti-lipid peroxidation, and magnesium deficiency leads to the increase of lipid peroxidation, also, magnesium deficient animals are more sensitive to oxidative stress than normal animals.

## Conclusions

In conclusion, among the three popular drinking waters, bottled natural water had suitable pH level, is the richest in minerals especially calcium and magnesium, and showed the best benefit for the development, lipid metabolism and antioxidant capacity of juvenile zebrafish. Tap water also showed good results, while bottled purified water were not suitable for juvenile zebrafish due to the deficiency of calcium and magnesium. Abundance of minerals is important for drinking water, so our results may have important implications for consumers especially adolescents who are in the important period of growth and development to select proper drinking water.

## Acknowledgements

The authors thank Key Laboratory for Molecular Animal Nutrition, Ministry of Education, College of Animal Sciences, Zhejiang University for animal care and maintenance.

## Author contributions

Conceived and designed the experiments: Jiajin Zhu and Cheng Chang. Performed the experiments: Cheng Chang. Performed lab analyses: Cheng Chang. Analyzed the data: Cheng Chang. Wrote the paper: Cheng Chang. All authors read and approved the final manuscript.

## List of symbols and abbreviations

dpf: days post-fertilization
TG: triglyceride
TC: cholesterol
LDL-C: low-density lipoprotein cholesterol
HDL-C: high-density lipoprotein cholesterol
SOD: superoxide dismutase
MDA: malondialdehyde
AI: Adequate Intakes
DBPs: disinfection by-products
TDS: total dissolved solids
TWG: fish water group
PWG: purified water group
NWG: natural water group
TWG: tap water group
RCT: reverse cholesterol transport

